# Precise Cas9 targeting enables genomic mutation prevention

**DOI:** 10.1101/058974

**Authors:** Alejandro Chavez, Benjamin W Pruitt, Marcelle Tuttle, Rebecca S Shapiro, Ryan J Cecchi, Jordan Winston, Brian M Turczyk, Michael Tung, James J Collins, George M Church

**Affiliations:** Wyss Institute for Biologically Inspired Engineering, Harvard University, Cambridge, Massachusetts, USA.; Department of Pathology, Massachusetts General Hospital, Boston, Massachusetts, USA.; Department of Genetics, Harvard Medical School, Boston, Massachusetts, USA.; Institute for Medical Engineering & Science, Massachusetts Institute of Technology, Cambridge, Massachusetts, USA.; Synthetic Biology Center, Massachusetts Institute of Technology, Cambridge, Massachusetts, USA.; ^6^Department of Biological Engineering, Massachusetts Institute of Technology, Cambridge, Massachusetts, USA.; Broad Institute of MIT and Harvard, Cambridge, Massachusetts, USA.

## Abstract

Here we present a generalized method of guide RNA “tuning” that enables Cas9 to discriminate between two target sites that differ by a single nucleotide polymorphism. We employ our methodology to generate a novel *in vivo* mutation prevention system in which Cas9 actively restricts the occurrence of undesired gain-of-function mutations within a population of engineered organisms. We further demonstrate that the system is scalable to a multitude of targets and that the general tuning and prevention concepts are portable across engineered Cas9 variants and Cas9 orthologs. Finally, we show that the designed mutation prevention system maintains robust activity even when placed within the complex environment of the mouse gastrointestinal tract.

---

There are few biological perturbations that rival point mutations when it comes to the power to affect phenotypic change. Single-base substitutions endow pathogens with resistance to antibiotics and rogue cells with oncogenic potential^1,2^. Despite decades of research into the causes and effects of point mutations, no tool exists to directly prevent their occurrence. At best, we can track when mutations occur and utilize them as prognostic factors to predict emergent properties or chemotherapeutic outcome^3,4^. Here we describe an *in vivo* “mutation prevention” system that can prevent the occurrence of targeted point mutations with little to no latent toxicity to the host organism.

The system is predicated on *Streptococcus pyogenes* Cas9 (hereon referred to as Cas9) and its orthologs: endonucleases directed to a target locus via hybridization between an associated guide RNA (gRNA) and a target site near a required protospacer adjacent motif (PAM)^5–8^. While incredibly plastic, Cas9 suffers from difficult-to-predict non-specific activity, which can be tolerant to multiple mismatches between the gRNA and the inappropriately bound locus^9–20^. Several methods exist to increase Cas9 specificity; however, no method has been proven to reliably enable the single-nucleotide resolution necessary for mutation prevention^13,21–28^. In order to endow Cas9 with the ability to discriminate between single-nucleotide polymorphisms (SNPs), we developed a screening methodology that confers single-nucleotide specificity through the selection of a tuned guide RNA (tgRNA).

To demonstrate the feasibility of preventing the emergence of point mutations with a Cas9/tgRNA system, we generated a strain of E. coli deficient in mismatch repair (MG1655-*mutS::kan*), to increase mutation rates, and harboring a plasmid encoding a catalytically-inactive version of the TEM-1 β-lactamase (TEM-1-S68N)^29,30^. The active version of TEM-1 confers resistance to β-lactam antibiotics, such as ampicillin^31^. Under typical growth conditions, errors in DNA replication and repair lead to the accumulation of mutations over time. Some of these mutations result in the reversion of the inactive N68 allele to its catalytically active form, S68. The number of revertants within the population is quantified by plating overnight cultures to solid media containing ampicillin (Fig. 1A).

We postulated that we could prevent these reversions by introducing a Cas9 system tuned to cut the active TEM-1 variant while exhibiting undetectable activity against the inactive allele. In practice, such a system would benignly persist within cells until the occurrence of the reversion. Cas9 would then cut at the active allele, causing irreparable genetic damage and thus a decrease in the overall number of ampicillin-resistant cells (Fig. 1B). By design, the inactive form of TEM-1 differs from the active form by a single nucleotide substitution (203G>A). Our initial, naïve system included a gRNA with full complementarity to the active S68 allele (203G) and thus only differed from the inactive N68 allele by a single nucleotide. As expected, this approach performed poorly, as the Cas9/gRNA system cut both the active and inactive forms efficiently. Consequently, the system exhibited high toxicity even in the absence of the reversion (data not shown). Furthermore, we found that even the high fidelity eCas9 and Cas9-HF1 variants failed to confer single nucleotide resolution with the naive one-off gRNA (**Supplementary Fig. 1**)

**Figure 1.**
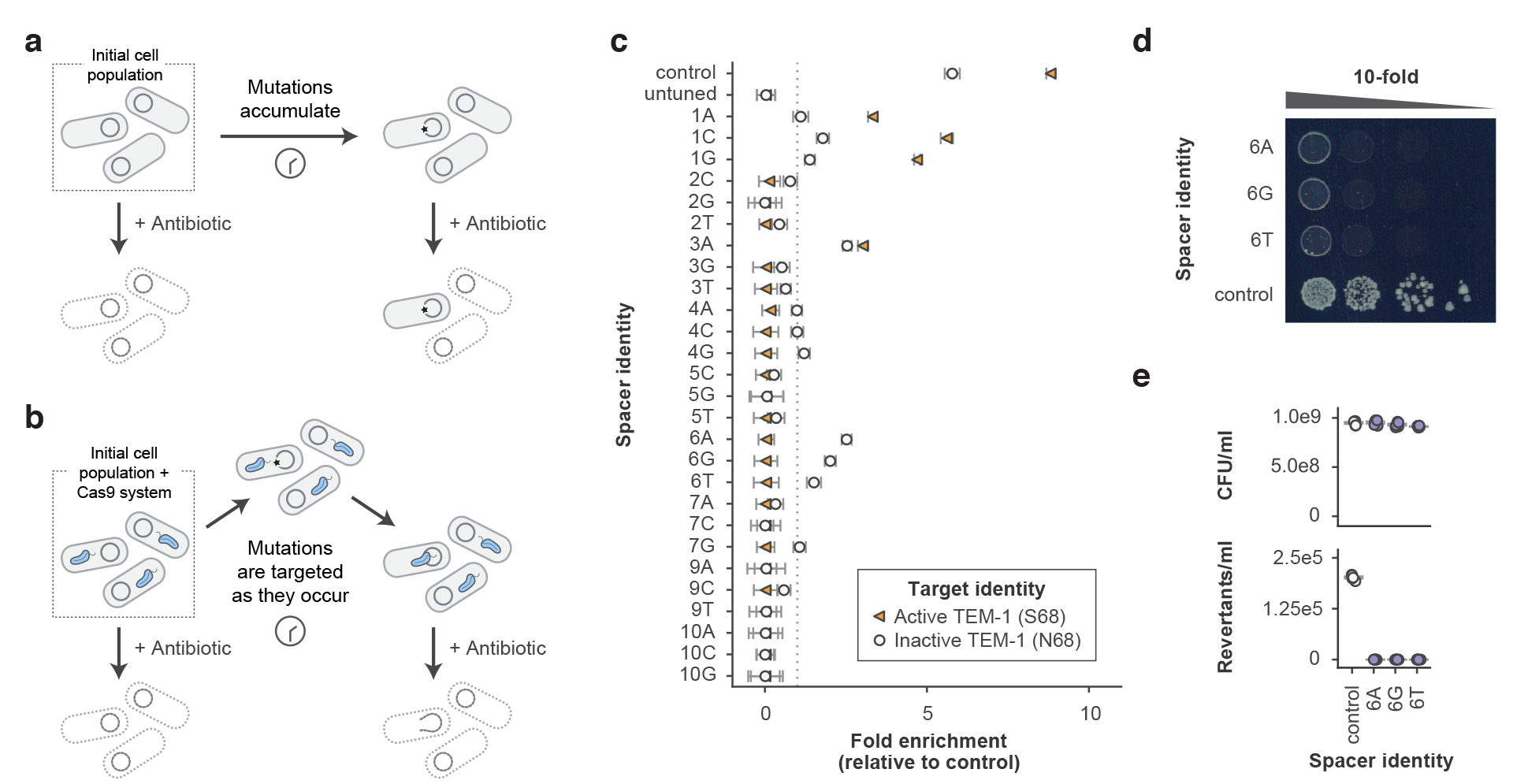
Mutation prevention system overview and performance. (a) Mutations conferring antibiotic resistance occur stochastically over time in native bacterial populations. When antibiotic pressure is applied to the population, cells with resistance-conferring mutations survive, while wild-type cells die (b) Cells with the mutation prevention system grow and divide normally. When a targeted resistance-causing mutation occurs it is cut by Cas9, causing its loss from the population. (c) Screening for tgRNAs designed to prevent the reversion of inactive TEM-1 to its catalytically-active form (conferring ampicillin resistance). In this assay, properly tuned gRNAs are expected to be enriched in the presence of the inactive target (N68) and depleted in the presence of the active target (S68). The y-axis indicates the additional tuning mismatch that is inserted into each respective gRNA. The dotted line represents a relative fold enrichment of one, above which library members are considered enriched and below which they are considered depleted; *n=3*, independent biological replicates. (d) Representative spot assay demonstrating the performance of mutation prevention systems containing the three most discriminatory tgRNAs (as indicated by the screening process). Each spot represents a 10-fold serial dilution. (e) The most discriminatory tgRNAs consistently prevent mutations to baseline levels, while not affecting the overall number of colony forming units present within the culture under non-selective conditions; *n=5*, independent biological replicates. All reversion rates are significant relative to the control (*P<0.01*), all CFU/ml counts are not significant relative to the control (*P>0.01*). For all experiments the control gRNA represents a guide that targets a sequence not present within the *E. coli* genome. All error bars represent S.E.M.

Previous work examining the factors that influence Cas9 behavior has shown that its activity is easily modulated by the introduction of mismatches within the gRNA that prevent full complementarity between it and the target site^51011^. We posited that, for a given pair of targets that differ by a single base, there may exist a set of mismatches within the gRNA that would eliminate activity on one of the variants while maintaining robust activity on the other. To test our hypothesis, we screened a library of gRNAs that each differed from the active TEM-1 allele (S68) by a single mismatch and from the inactive TEM-1 allele (N68) by two mismatches (**Supplementary Fig. 2**). We focused our mutational analysis on the region within the gRNA that interacts with the bases proximal to the PAM at the target locus, as these residues have been shown to be most critical for Cas9 activity^5,9,10,11^.

Our screen was designed such that an ideal tgRNA candidate was expected to be markedly depleted when in the presence of the functional S68 allele (signifying that it cut the undesired TEM-1 variant that we wish to prevent) and strongly enriched when tested against the non-functional N68 allele (suggesting that it did not cut the desired TEM-1 variant we wish to maintain). Upon performing the screen, we identified several promising tgRNA candidates that were subjected to further characterization (Fig. 1C).

Individual testing of these select library members revealed that the fold depletion we observe within the screen strongly correlates with their respective activities in isolation (**Supplementary Fig. 3**). This validates the capability of our library approach to accurately identify the most discriminatory gRNAs. Based on the results of ourscreen and subsequent validation, we selected the three candidate tgRNAs with the greatest discriminatory power, 6A, 6G, and 6T, for additional analysis. We tested their ability to prevent emergence of the active TEM-1 allele from within a population of cells containing only the inactive TEM-1 variant. As predicted, all three tgRNAs were able to prevent reversion to the active TEM-1 allele by several orders of magnitude over cells with a non-functional control gRNA, while also exhibiting minimal levels of baseline toxicity (Fig. 1D,E).

Because Cas9 represents one of thousands of known Cas9 proteins, each with differences in efficiency, specificity, and PAM requirements, we sought to verify that our approach was generalizable to two commonly used Cas9 orthologs: NM-Cas9 and ST1-Cas9^32–33^. We leveraged the same TEM-1 reversion assay and gRNA screening technique to identify tgRNAs with high discriminatory power between the active and inactive targets. Each screen yielded promising tgRNA candidates that proved to be highly effective at preventing the TEM-1 reversion mutation with either NM-Cas9 or ST1-Cas9, respectively (**Supplementary Fig. 4**).

Having demonstrated the efficacy of the mutation prevention concept with our exogenous TEM-1 model, we sought to apply the same approach to prevent endogenous mutations. Toward this goal, we set out to prevent several common mutations within *E. coli* that confer resistance to two clinically-relevant antibiotics: streptomycin and rifampicin^34^.

Streptomycin is an aminoglycoside antibiotic that inhibits protein synthesis by binding to bacterial ribosomes^35^. There are well-characterized ribosomal protein mutations that confer resistance to streptomycin, including commonly observed substitutions in *rpsL^36^.* We designed mutation prevention systems for two of the highest frequency resistance-bearing mutations within *rpsL:* 128A>G and263A>G. Our screening process yielded two effective tgRNAs, both of which prevented the occurrence of their targeted mutations to below the detection limits of our next-generation sequencing assay, while causing no apparent toxicity to non-mutant cells (**Fig. 2A and Supplementary Tables 1,2**).

**Figure 2.**
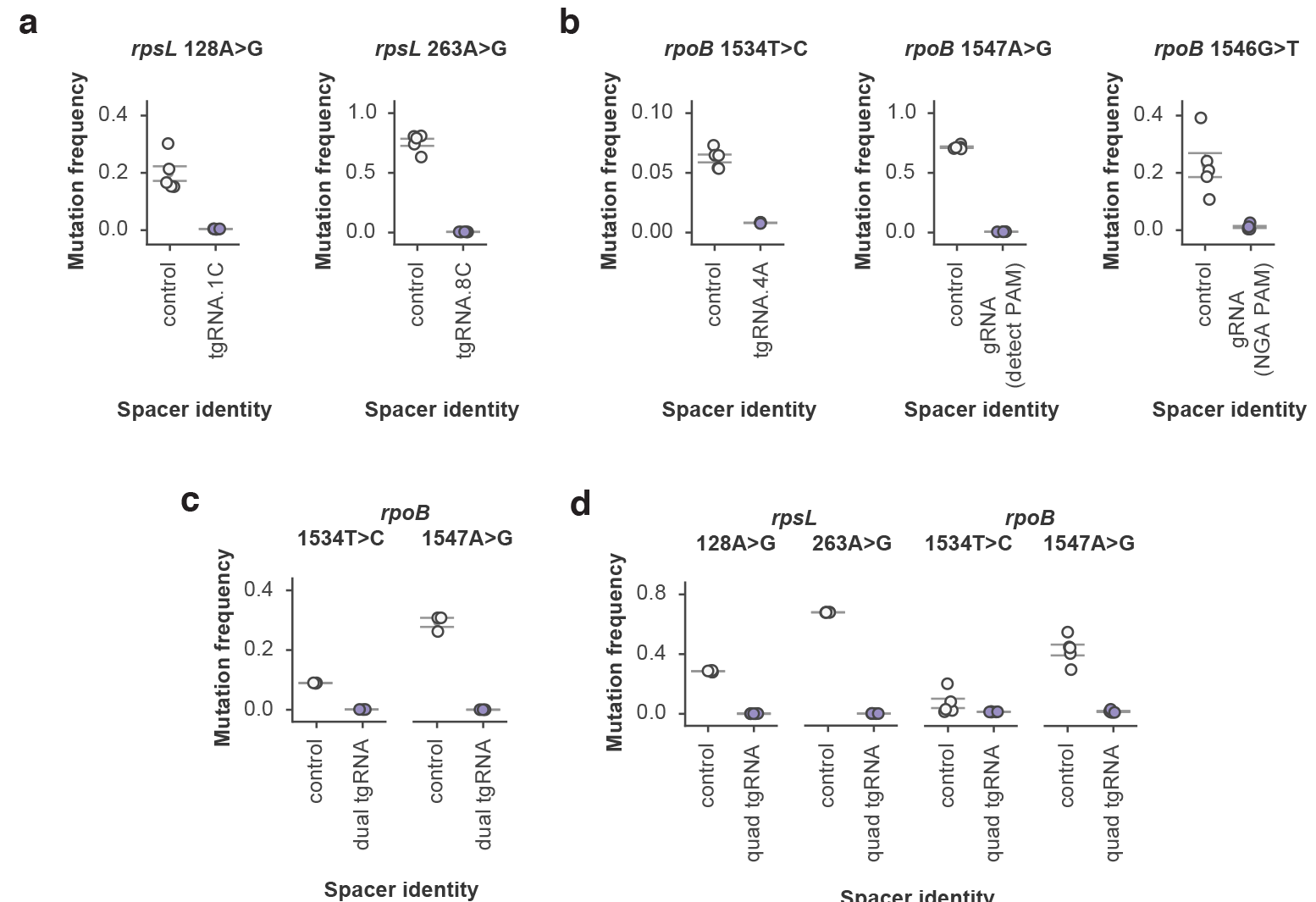
Endogenous mutation prevention within E. coli. (a) Frequency of rpsL 128A>G and rpsL 263A>G alleles within populations of streptomycin-resistant cells containing either tgRNA rpsL.128A>G.1C or tgRNA rpsL.263A>G.8C, respectively, as compared to cells with a control gRNA. (b) Frequency of rpoB 1534T>C, rpoB 1547A>G and rpoB 1546G>T mutations within populations of rifampicin-resistant cells. For rpoB 1534T>C, a tgRNA containing an additional mismatch at the-4A position was employed to prevent the occurrence of the mutation. With rpoB.1547A>G, the mutation generates a novel PAM that induces Cas9 targeting. Because the rpoB.1546G>T mutation is not near a conventional Cas9 PAM, the Cas9-VQR variant with an alternative PAM requirement was employed along with an appropriate gRNA in order to prevent the mutation. (c) Simultaneous multiplex mutation prevention was performed against both rpoB 1534T>C and 1547A>G mutations in the same population of rifampicin-resistant cells by simultaneously expressing a pair of tgRNAs (denoted as dual tgRNA) along with the necessary Cas9 machinery. (d) Similar to panel C except a total of four mutations (rpsL 128A>G and 263A>G, rpoB 1534T>C and 1547A>G) were simultaneously prevented by using an array of four previously validated tgRNAs (denoted “quad tgRNA”) along with Cas9 protein. For all experiments the control gRNA represents a guide that targets a sequence not present within the E. coli genome. All error bars represent S.E.M., n = 5 independent biological replicates for all experiments, except for subpanel c (n = 3). Experimental means are significant relative to control means (P < 0.01) with the exception of rpob.1534T>C.4A in subpanel d (P = 0.14).

Rifampicin is a widely used antibiotic that inhibits RNA polymerase function, with resistance arising from several well documented mutations within the *rpoB* gene^37,38^. To further demonstrate the plasticity of our approach, we targeted a series of the prominent rifampicin-resistance mutations and generated unique tgRNAs against each using three disparate design principles. To prevent the *rpoB* 1534T>C mutation, we screened a series of gRNAs and identified a canonical tgRNA, bearing additional mutations that was highly efficient (**Fig. 2B**). Targeting the *rpoB* 1547A>G mutation was simplified by the fact that the mutation generates a PAM that is not present within the wild-type *rpoB* gene. A gRNA designed to bind in the presence of the generated PAM conferred marked mutation prevention (**Fig. 2B**). Finally, to prevent the *rpoB* 1546G>T mutation, we employed the Cas9-VQR variant with altered PAM specificity^39^. This was necessary given the lack of a canonical PAM sequence near the mutation site. Cas9-VQR, in the presence of an appropriate gRNA, led to efficient mutation prevention (**Fig. 2B**).

Having demonstrated both the portability and flexibility of our mutation prevention concept, we next assessed its scalability by targeting multiple mutations within *rpoB* with a single system. There was no obvious decrease in efficiency upon multiplexing, with both *rpoB* 1534T>C and 1547A>G mutations being simultaneously prevented within the engineered strain (**Fig. 2C**). To further probe the limits of our multi-mutation prevention system we sought to simultaneously inhibit all of the previously targeted endogenous mutations (except *rpoB 1546G>T,* which requires the Cas9-VQR variant) within a single strain. Cells carrying a full complement of four tgRNAs exhibited minimal toxicity as compared to unmutated cells and retained robust mutation prevention against all targeted loci (**Fig. 2D and Supplementary Table 1**).

We next wanted to assess the robustness of our system by preventing point mutations from emerging within a complex *in vivo* environment. Towards this goal we inoculated gnotobiotic mice with one of two strains containing either a pair of non-targeted gRNAs or a pair of tgRNAs directed to prevent mutations that confer rifampicin resistance within *E. coli.* Two days after colonization, mice were provided rifampicin within their drinking water and the spectrum of rifampicin resistance mutations was analyzed in bacterial cells recovered from fecal samples collected on days 4 to 7, post-colonization (**Fig. 3A**). We observed minimal frequencies of the targeted point mutations within the engineered strain, relative to the much higher frequencies within the control (**Fig. 3B**).

**Figure 3.**
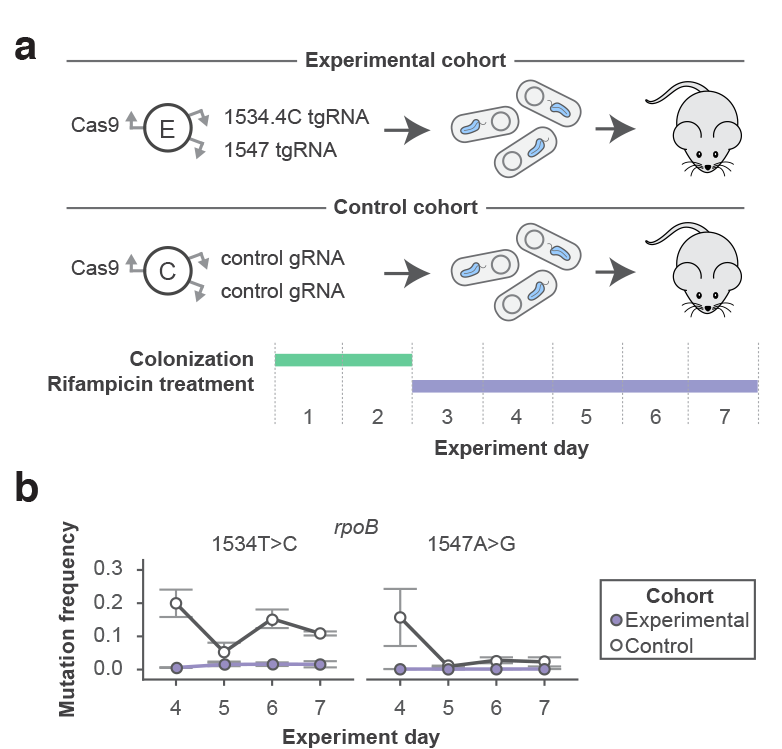
Longitudinal mutation prevention in a complex environment. (a) Two cohorts of mice were colonized with either an experimental *E. coli* strain encoding the dual *rpoB* 1534T>C / 1547A>G mutation prevention system or a control *E. coli* strain encoding a system with two gRNAs that do not cut anywhere in its genome. After a two day initial colonization, mice were provided water containing the antibiotic rifampicin for the remaining five days of the experiment. (b) The mutation prevention system successfully prevented the occurrence of the two targeted mutations for the duration of the rifampicin treatment. All error bars represent S.E.M., *n;=3* experimental mice and *n=2* control mice. The difference between experimental and control frequencies in panel b is significant days 4, 6, and 7 for 1534T>C and day 4 for 1547A>G (P<0.01).

It is expected that the mutator strain of *E. coli* utilized within all of our experiments would commensurately increase the frequency of mutations that disable the mutation prevention system itself^3,29^. Given that the system is likely breaking at an appreciable rate, the fact that we do not see obvious degradation in its performance over time likely indicates that the system places a negligible fitness burden on the host organism. If the system were to place a significant fitness burden on its host, one would expect individuals with attenuated or broken systems to rapidly overtake the population, thus facilitating escape from its selection. It is thanks to this minimal fitness burden that escape is an extremely rare event. This is due to the fact that during any given point in time there is only a small population of cells within the total population that have inactivated the system and it is within this rare population that a second targeted gain-of-function mutation must occur. This “two hit” requirement effectively makes escape from our mutation prevention strategy an extremely rare event and explains our efficacy at preventing undesired mutations. The robust performance of the mutation prevention system within the mouse gastrointestinal tract is particularly notable given its tolerance of multi-day rifampicin selection, the absence of the active selection for the episomal plasmid encoding our system, and the fact that these experiments were performed within a highly mutagenic mismatch repair deficient background of *E. coli.*

Although our mutation prevention strategy requires that users know the identity of the mutations that they wish to prevent, we note that there is a plethora of previously characterized gain-of-function mutations that endow living cells with undesired phenotypes. In these cases, users may want to prevent a subset of mutations in order to gain deeper insight into the existence and effects of other rare alleles. In addition, thanks to the rapid increase in the throughput and adoption of next generation DNA sequencing, we stand on the cusp of what will be the eventual systematic characterization the various genomic mutations that enable cells to evade the selective pressures imposed upon them by both natural and artificial systems. We forsee future applications of our method in enabling users to capitalize upon this expansive characterization, and in turn, manipulate it to prevent undesired outcomes (while also gaining even deeper understanding of evolutionary processes).

The marked efficiency of the described mutation prevention system makes it an appealing platform for use in a variety of applications — from the study of evolutionary adaptation under stress to the creation of novel forms of biocontainment, in which the target upon which selection occurs is at the single nucleotide level^40,41^. We further expect that the generalizable principle of guide RNA tuning will function synergistically with newly engineered, high-specificity variants of Cas9^26,27^, as well as other RNA/DNA guided endonucleases like Cpf1, NgAgo, and C2C 2^42–45^. As a result, applications requiring true single-nucleotide specificity, such as the disabling of diseased alleles delineated by SNPs, will finally be realized.

Along with its immediate applications, we foresee longer term implications of our work in paving the way towards the creation of “genomic immune systems”, that prevent undesired genetic alterations (e.g., gain-of-function mutations) such as oncogenic mutations within engineered organisms^2^. In addition, the mutation prevention systems we describe here might one day be combined with gene drives to shape mutational landscapes within wild populations and by doing so enable science to hinder the spread of undesired traits, such as herbicide resistance among invasive plants^46,47^.

## Acknowledgements

We would like to thank L. Certain, X. Guo, A. Tsou, A. Tung, S. Vora, N.C. Yeo and J. Zhu for helpful discussions and technical assistance. We acknowledge research support from the US National Institutes of Health National Human Genome Research Institute grant P50 HG005550, and the Wyss Institute for Biologically Inspired Engineering. A.C.was funded by the National Cancer Institute grant 5T32CA009216-34. J.J.C. was supported by the Defense Threat Reduction Agency grant HDTRA1-15-1-0051.

### Contributions

A.C. conceived of the mutation prevention concept. B.W.P. conceived of the gRNA screening pipeline. A.C. and B.W.P. conceived of and designed experiments. A.C, B.W.P., M. Tuttle, R.S.S., R.J.C., B.M.T., M. Tung, and J.W. performed experiments. J.J.C. and G.M.C. supervised the study. B.W.P. and A.C. wrote the manuscript with support from all authors.

### Conflict of interest

G.M.C. is a founder and advisor for Editas Medicine. G.M.C. has equity in Editas, Enevolv and Caribou Biosciences (for full disclosure list, please see: <u>http://arep.med.harvard.edu/gmc/tech.html</u>). A provisional patent has been filed by Harvard University related to the work within this manuscript.

